# The temporal specificity of BOLD fMRI is systematically related to anatomical and vascular features of the human brain

**DOI:** 10.1101/2024.02.01.578428

**Authors:** Daniel E. P. Gomez, Jonathan R. Polimeni, Laura D. Lewis

## Abstract

The ability to detect fast responses with functional MRI depends on the speed of hemodynamic responses to neural activity, because hemodynamic responses act as a temporal low-pass filter which blurs rapid changes. However, the shape and timing of hemodynamic responses are highly variable across the brain and across stimuli. This heterogeneity of responses implies that the temporal specificity of fMRI signals, or the ability of fMRI to preserve fast information, could also vary substantially across the cortex. In this work we investigated how local differences in hemodynamic response timing affect the temporal specificity of fMRI. We used ultra-high field (7T) fMRI at high spatiotemporal resolution, studying the primary visual cortex (V1) as a model area for investigation. We used visual stimuli oscillating at slow and fast frequencies to probe the temporal specificity of individual voxels. As expected, we identified substantial variability in temporal specificity, with some voxels preserving their responses to fast neural activity more effectively than others. We investigated which voxels had the highest temporal specificity, and tested whether voxel timing was related to anatomical and vascular features. We found that low temporal specificity is only weakly explained by the presence of large veins or cerebral cortical depth. Notably, however, temporal specificity depended strongly on a voxel’s position along the anterior-posterior anatomical axis of V1, with voxels within the calcarine sulcus being capable of preserving close to 25% of their amplitude as the frequency of stimulation increased from 0.05Hz to 0.20Hz, and voxels nearest to the occipital pole preserving less than 18%. These results indicate that detection biases in high-resolution fMRI will depend on the anatomical and vascular features of the area being imaged, and that these biases will differ depending on the timing of the underlying neuronal activity. While we attribute this variance primarily to hemodynamic effects, neuronal nonlinearities may also influence response timing. Importantly, this spatial heterogeneity of temporal specificity suggests that it could be exploited to achieve higher specificity in some locations, and that tailored data analysis strategies may help improve the detection and interpretation of fast fMRI responses.

## 1. Introduction

Blood-oxygenation-level-dependent (BOLD) functional MRI (fMRI) is the most commonly used imaging technique for the non-invasive study of cognitive processes in the human brain because of its submillimeter and subsecond resolution. Many ongoing advancements in fMRI techniques have increased the spatial resolution and sensitivity of BOLD fMRI (Blazejewska et al., 2019; Dumoulin et al., 2018; Mareyam et al., 2020; Polimeni et al., 2015; Viessmann and Polimeni, 2021 and other references therein). However, improving temporal resolution is often considered more challenging. The temporal constraints of BOLD functional imaging are ultimately determined by the inherent biological timescales of neurovascular coupling and the resulting hemodynamics (Buxton et al., 2014) Understanding the temporal precision of the BOLD response is therefore important for extracting precise timing information about the neuronal activity of interest from fMRI data.

Although the neuronal activity of interest often happens at millisecond scales, BOLD fMRI only measures the subsequent hemodynamic changes that unfold on the scale of seconds. Common expectations based on hemodynamic response models developed in the early days of fMRI (Glover, 1999) suggested that neurovascular coupling is slow, yielding BOLD responses expected to be below ∼0.15 Hz. However, resting-state fMRI studies have demonstrated meaningful spatial structure in high-frequency BOLD signals (Boubela et al., 2013; Chen and Glover, 2015). Furthermore, task-based fMRI studies have shown that the BOLD responses can track external stimuli with frequencies as high as 0.75 Hz (Lewis et al., 2016), demonstrating that hemodynamic responses to neuronal activity can be surprisingly faster than previously believed. Imaging fast BOLD responses could open a window to probe rapidly evolving neuronal activity, such as those associated with decision-making, language-processing, attention, and sensory-motor integration (Polimeni and Lewis, 2021). However, a deeper understanding of the temporal properties of hemodynamic responses is needed to analyze and interpret these fast signals.

A challenge for identifying precise timing information in fMRI is that the temporal properties of hemodynamic responses are highly heterogeneous across the brain (Bailes et al., 2023; Handwerker et al., 2004; Pfeuffer et al., 2003) and across experimental contexts (Chen et al., 2021; Friston et al., 1998; Handwerker et al., 2012). BOLD responses are influenced by the vascular architecture of the cortex: they are weaker but have an earlier onset in the parenchyma, and stronger in the large draining veins that collect large volumes of deoxygenated blood from multiple capillary beds (Turner, 2002). Since the vascular architecture is organized such that veins drain upwards towards the pial surface (Duvernoy et al., 1981), this effect is not only visible when comparing parenchyma and venous responses (Gati et al., 1997; Kay et al., 2019; Lai et al., 1993; Siero et al., 2009; Uludağ and Blinder, 2018; de Zwart et al., 2005), but by proxy also when comparing responses across cortical depths (Siero et al., 2014; Siero et al., 2011) given that responses at the pial surface are more likely to include larger veins. Since hemodynamic properties act as a filter on the underlying neuronal activity, they thus determine the timing and amplitude of measurable BOLD responses: areas with slower, sluggish hemodynamic responses should produce correspondingly slow signals, whereas sharper and faster responses should be able to robustly track high-frequency signals. The intrinsic temporal resolution of fMRI should therefore vary across the brain depending on local hemodynamic timing.

In addition to modulating the temporal precision of fMRI, this vascular heterogeneity can introduce detection biases (Polimeni et al., 2018) in BOLD responses that become increasingly pronounced at higher spatial and temporal resolution imaging. These biases can have profound effects depending on an fMRI study’s experimental design. For studies with slow task designs, small timing differences can sometimes be negligible since they can be accounted for during statistical analysis—using a general linear model (GLM) for example—by modeling not only the response predicted by a convolution with a standard hemodynamic response function (HRF), but also by modeling its temporal derivatives, or using flexible modeling approaches. However, for rapid task designs (e.g. event-related or naturalistic stimulus designs), these biases can lead to several issues, such as undetected responses (false negatives), spatial errors when creating topographic maps; and misinterpretation of the BOLD signal when comparing responses to stimuli with different timings. Any biases that are imparted by vascular anatomy will likely impact distinct cortical regions differently (Handwerker et al., 2004), and indeed differences in hemodynamic response delays on the order of seconds and large response amplitude differences have also been observed (Amemiya et al., 2020).

Here, we investigated how the temporal precision of BOLD fMRI signals varies across the human visual cortex, to test whether vascular or other anatomical features may determine, in part, where fast responses can be detected. We used ultra-high field fMRI (7 Tesla) at high spatiotemporal resolution (1.06 mm isotropic resolution with 0.874 s temporal resolution) to measure responses to rapidly varying visual stimuli across cortical depths and close to large veins. Investigating these fast responses at high spatiotemporal resolution requires careful experimental considerations: high-frequency (> 0.2 Hz) responses to rapidly fluctuating stimuli are typically too small in amplitude to measure at the single-voxel level, demanding an impractical amount of trial-averaging for reliable detection above the noise floor, hindering a direct analysis of local response properties at the single-voxel level. For this reason, we used 0.20 Hz as our fastest stimulus frequency and analyzed the frequency response of each voxel to infer how well it can preserve fast information. We defined a metric for “temporal specificity”, by calculating the amplitude at faster frequencies relative to the amplitude at a slower reference frequency (in this study, 0.05 Hz). This “temporal specificity” metric represents how well a voxel preserves information about responses to fast stimuli (as compared to slow stimuli). For sharp HRFs, the amplitude at fast frequencies is expected to be higher, since sharp HRFs preserve more high-frequency information. Based on prior work on the relationship between responses and vascular-anatomical features (Siero et al., 2011; de Zwart et al., 2005), we hypothesized that deep layers of cortical gray matter would have increased temporal specificity since they have earlier responses, and for the same reason parenchyma would have increased specificity when compared to large veins. We found that oscillatory stimulation elicited strong single-voxel BOLD responses across all frequencies tested, allowing us to robustly stratify responses according to cortical depth, vascular compartment, and anatomical position along V1. Our results demonstrated that temporal specificity differences were weakly related to cortical depth and vascular compartment yet varied strongly along the anterior-posterior axis of V1. Importantly, our results indicate that fast BOLD responses will be more readily detectable in regions with higher temporal specificity, which is spatially structured and linked to brain anatomy.

## 2. Methods

### 2.1. Subject population

All subjects provided informed written consent, and all procedures were approved by Massachusetts General Hospital’s Institutional Review Board. A total of 16 subjects with previous MRI experience were recruited and scanned, and data from 12 subjects (age 26.6 *±* 3.2 years, 6F/6M) were analyzed. Subjects were excluded if motion was larger than 2.5 mm throughout the session, or if the behavioral performance on the behavior task (explained below) in any run was found below 75%. Three subjects were excluded for motion and one for not completing the full experiment.

### 2.2. MRI Data Acquisition

Participants were scanned on a 7 T Siemens (Magnetom “Classic”, Erlangen, Germany) whole-body scanner with a custom-built 32-channel head coil array and birdcage head coil for transmit. Each session began with a 0.75-mm isotropic dual-echo MPRAGE (Van der Kouwe et al., 2008) as an anatomical reference with the following parameter values: TR = 2530 ms, TI = 1100 ms, TE = 1.74 and 3.7 ms, flip angle = 7*^◦^*, outer-loop acceleration = 2 and no partial Fourier, bandwidth=651 Hz/px, and a duration of 7:20 minutes. After the anatomical reference scan, eleven BOLD functional runs were acquired using a single-band single-echo gradient-echo 2D EPI protocol. Coronal slices were positioned over the occipital pole targeting the calcarine sulcus. The protocol had following parameter values: 16 slices with 1.06-mm isotropic resolution (R = 4 in-plane acceleration and no partial Fourier), TR = 874 ms, TE = 24 ms, flip angle = 52*^◦^*, matrix size = 164 *×* 164, bandwidth = 1452 Hz/px, nominal echo spacing 0.81 ms, phase-encoding direction = Left *→* Right. Each functional run collected 300 volumes for a total acquisition time of 257 s. Between the 4^th^ and 5^th^ run a larger-FOV EPI protocol with 40 slices of same resolution was acquired to aid the registration between the small-FOV BOLD data and the full-brain anatomical scan. Between the 7^th^ and 8^th^ run a 3-minute inversion-recovery EPI (Renvall et al., 2016) was acquired, but not used in this analysis. For quality assessment purposes, temporal signal-to-noise (tSNR) maps were computed from a resting-state run acquired in one subject using the same protocol used for our visual stimulation data (Supplementary Figure 1). Mean tSNR values within the same ROI used in our functional data analysis were greater than 10, as expected based on our voxel size (see e.g. (Triantafyllou et al., 2011)).

### 2.3. Visual stimulus

Visual stimuli were presented using a DLP projector (Psychology Software Tools), with timing synchronized to the 60Hz refresh rate of the stimulus delivery computer, onto a screen placed at the back of the scanner bore. Subjects viewed the stimulus through an angled mirror placed above their eyes positioned 110 cm away from a projector screen measuring 40 cm by 30 cm. The field-of-view of participants thus covered approximately 22° (horizontal) by 16° (vertical) of visual field. Stimulus presentation code was written in Lua using the Löve 2D framework (https://love2d.org), and timing accuracy was checked against a 120Hz iPhone 12 camera by filming the presentation outside of the scanner as it would be displayed in the scanner and comparing its timing to the experimental design. The visual stimuli consisted of 12Hz counterphase flickering radial “checkerboards” (radially spaced at every 18°yielding 20 sectors, and annually spaced by sin(10 × log(radius × radius))), thresholded at 0, presented continuously, beginning 14 s after the start of the first volume of the scan. The luminance contrast of the checkerboards was sinusoidally modulated from 0 (no contrast) to 0.8 (where 1 would be maximum black/white contrast) at frequency of interest throughout the runs as previously described (Lewis et al., 2016). An implementation of this task as used in the current experiments can be found under https://github.com/dangom/love-experiments. For the experiment, eleven functional runs were acquired: 2 with luminance modulated at 0.05 Hz, the first of which was used as a functional localizer, 3 modulated at 0.10 Hz, and 6 modulated at 0.20 Hz. Each trial was defined as a full cycle of the sine period. The order of the stimuli was set to [0.05, 0.20, 0.10, 0.20, 0.20, 0.10, 0.20, 0.05, 0.20, 0.10, 0.20] Hz to avoid order effects. Multiple runs were acquired for averaging. During visual stimulation runs, subjects performed a simple visual fixation task: a red dot at the center of the screen alternated between light and dark red with switch times drawn from a uniform distribution between 0.8 and 3 s; subjects were instructed to press a button on an MR-compatible USB button box every time the dot changed color.

### 2.4. Data Preprocessing

#### 2.4.1. Anatomical preprocessing

As described in (Van der Kouwe et al., 2008), multiple echoes of the MP-RAGE acquisition were combined using root sum of squares before any further processing.

##### 1. Bias field correction

Anatomical images were then bias field corrected using the joint bias-field and class segmentation estimation provided by SPM 12 (https://www.fil.ion.ucl.ac.uk/spm/software/spm12/). The bias field correction requires setting two parameters: the smoothness of the bias-field, and the bias regularization. These parameter values have a strong impact on the trade-off between estimating the bias field or the segmented tissue classes. Because we were interested in the portion of the visual cortex covered by the 16-mm FOV where contrast between gray and white matter is subtle, we chose conservative parameters that penalize large variations in the bias field (biasreg = 0.15, biasfwhm = 20). This choice opts to underestimate the true bias field but reduces the chance of inadvertently misclassifying the GM/WM intensity contrast as local bias field. These values were chosen after visual inspection of the bias field and reconstructed cortical surfaces when preprocessing data.

##### 2. Cerebral cortical surface reconstruction

After bias-field correction, cortical surfaces were automatically reconstructed using FreeSurfer 7.1.1 (https://surfer.nmr.mgh.harvard.edu). Because V1 intensities at 7T are higher than average brain gray matter values, recon-all was run with flags -seg-wmlo and -seg-grayhi set to 95 and 105, respectively. This reduces the likelihood of surfaces being incorrectly placed in the middle of the GM ribbon. The brain mask generated by recon-all was also refined by removing areas which the SPM bias-field and segmentation algorithm had already classified as belonging to bone or other tissues. This was done after noticing that the automatically generated brain mask was at times too relaxed and did not fully remove the skull nor sagittal sinus.

#### 2.4.2. Functional preprocessing

##### 1. Slice time and motion correction

Slice time correction was performed using filtershift, interpolating the data to the middle slice of each volume (Parker et al., 2017). Motion correction was performed subsequently using AFNI’s 3dVolReg via FreeSurfer’s mc-afni2 wrapper.

##### 2. Within-subject inter-run coregistration and run averaging

All functional runs were co-registered to each subject’s functional localizer run. Transformations were estimated using ANTs (https://github.com/ANTsX/ANTs) in a hierarchical approach whereby first a rigid transform was estimated and used as input to estimate a similarity (rigid + scaling) transform. This was in turn used to estimate an affine transform which was used to co-register runs. (This approach is customary and recommended within the ANTs documentation to improve accuracy.) Co-registered runs were visually inspected, and runs of the same stimulus frequency were averaged using fslmaths.

##### 3. Registration to anatomical reference

A transformation between the functional localizer space and the anatomical space was computed to assign cortical depth estimates and anatomical ROI labels to each voxel in the functional run, using information from the FreeSurfer cortical surface reconstruction and parcellation in anatomical space. A transformation between the functional localizer space and the anatomical space was computed in two steps. The first step was identical to the inter-run coregistration using ANTs, but registers the localizer run to the larger-FOV reference EPI (functional *↔* large FOV). The second step co-registers the larger-FOV reference to the anatomical data using FreeSurfer’s boundary-based registration (larger-FOV *↔* anatomical) (Greve and Fischl, 2009). These two transforms were then concatenated to generate a mapping from functional localizer space to anatomical space (functional *↔* anatomical).

##### 4. Voxelwise cortical depth estimates

Once a transformation between functional and anatomical space had been established, it was possible to obtain the distance from the centroid of each voxel in functional space to the pial and white matter surfaces computed in anatomical space. Depth values are given in millimeters from the centroid of a voxel to the white matter surface (Polimeni et al., 2018, 2010). A value of 0mm represents a voxel exactly at the WM/GM interface, and higher values represent distances upwards into the GM and CSF. Because the cortical thickness varies across V1, however, a voxel with a depth of, e.g., 1 mm may be—depending on the thickness—either closer to the white matter or to the pial surface. In general, given the median cortical thickness of V1 being close to 2 mm, voxels with absolute depths between 0.7 and 1.3 mm should contain mostly gray matter, with voxels with depths below or above containing more partial volume of WM or CSF, respectively. (About 85.7 percent of voxels in the ROI used for analysis are contained mostly within the cortical gray matter. A partial volume analysis is shown in Supplementary Figure 2.)

##### 5. Laminar smoothing

Finally, because of the small amplitude of oscillatory responses in the parenchyma, especially at the 0.20Hz condition, a small amount of “laminar” smoothing (Blazejewska et al., 2019) was performed using LAYNII’s LAYERSMOOTH (Huber et al., 2021). For this, the gray matter ribbon was divided into 4 depths and smoothing was applied within voxels of the same depth using a gaussian filter with a full-width at half-max of 1.5mm.

### 2.5. Data Analysis

#### 2.5.1. Defining a localizer ROI to estimate oscillatory responses

A localizer ROI was defined for each subject based on both anatomical and functional considerations. The mask was created based on the following criteria: (1.) only voxels located within anatomically defined V1; and (2.) above-threshold positive activation in the functional localizer run. For this, first a V1 anatomical mask was created by registering the V1 prediction (i.e., the surface based V1_exvivo label) from FreeSurfer to each subject’s functional space. The mask resampling was performed with antsApplyTransforms using ITK’s GenericLabel interpolator with linear interpolation (Schaerer et al., 2014). Second, an activation mask was generated by thresholding Z (Gaussianized F) statistics at Z > 3.5 from activation maps from the averaged 0.05Hz runs (this choice of threshold does not qualitatively impact results; see Supplementary Figure 3). Activation maps were obtained using FSL FEAT (https://fsl.fmrib.ox.ac.uk/fsl/fslwiki/FEAT) (Woolrich et al., 2001), and using as regressors a sine and cosine waveform with the frequency of the localizer run, 0.05 Hz, and performing an F-test on the response to both sinusoid regressors. The analysis was run after discarding the initial 60 volumes of the acquisition, to minimize not only T1 transient magnetization effects but also transient hemodynamic effects related to the onset of the continuous stimulation. A high-pass filter with a cutoff of 1/25 s was included in the GLM for localizer definition, but high-pass filtered data were not used in any further analysis to avoid biasing results towards any frequency (Simon and Buxton, 2015) (results did not qualitatively change when high-pass filtering was used). No additional regressors were added as nuisance components. The mask was constrained to only include positive BOLD responses: the delay of all voxels from the 0.05Hz localizer run was estimated (estimation described below) and a mask was defined from all voxels whose onset delays were within 1–6 seconds after the stimulus. This interval was chosen based on our estimates of delays from conventional HRFs which suggest that delays should be well within that range. Negative BOLD responses were expected to be out of phase and thus have delays outside of that window. The localizer ROI mask was thus defined as the intersection of the anatomically defined V1 mask and the functionally-defined positive BOLD mask.

Finally, since more voxels were expected to be active at the pial surface of GM than in deeper GM due to the larger amplitude of pial-vein BOLD signals (Polimeni et al., 2010), the mask was further refined by only keeping voxels that were active at the pial surface that had a corresponding voxel active deep within the gray matter ribbon. To do that we used the cortical surface mesh vertex coordinates that represent cortical geometry: because FreeSurfer establishes a one-to-one vertex correspondence between the white and pial surfaces, it is straightforward to identify voxels intersecting the white matter surface that correspond to any given voxel intersecting the pial surface; this allowed voxels to be considered in pairs at both the inner and outer boundaries of the gray matter ribbon. All active voxels not intersecting the cortical ribbon (occasional false positives), or where only a single voxel of the pair was active, were removed to avoid an analysis biased towards surface voxels. Thus, the final functional ROI was created from the localizer ROI using this criterion.

#### 2.5.2. Estimating delay, amplitude and temporal specificity of oscillatory responses

To estimate delay and amplitude of the BOLD responses to visual stimulation, the time-series of each voxel in each averaged functional run (one per frequency per subject) was interpolated to a 100-ms grid using cubic interpolation. Subsequently percent signal changes were calculated within each voxel by subtracting the mean and dividing the result by the mean and multiplying that value by 100. Te mean was calculated over the steady-state response only (after discarding the first 54 seconds of each run to exclude transient effects). For comparison, computing the baseline as the first 10 seconds before stimulation led to an increase in the estimated PSC, but did not change our overall results (Supplementary Figure 4). All trials within each PSC-normalized time-series were then averaged to yield a cycle-average response for each voxel at each frequency for each subject. Finally, the amplitude of the oscillation was defined as the peak-to-trough magnitude of this cycle-averaged response. The delay was defined as the time of the trough of the response. The temporal specificity was defined as the ratio of the amplitude of responses at different stimulus frequencies. For example, the amplitude ratio “0.20 Hz/0.05 Hz” means the measured oscillation amplitude of a voxel to the 0.20Hz stimulus divided by the estimated amplitude to the 0.05Hz stimulus condition. This metric thus captured the ability of individual voxels to preserve their response magnitude to fast stimuli.

#### 2.5.3. Excluding false-positive voxels with a control analysis

As a control analysis to remove “false-positive” voxels that appeared to be responding to the stimulus but were doing so by chance, all data were re-analyzed as if the underlying stimulus had a frequency of 0.19 Hz. This resulted in a null distribution for amplitudes for each stimulus condition (0.05Hz, 0.10Hz and 0.20Hz). Voxels that had a true response amplitude at any frequency below the 50% of the null-distribution for that frequency were excluded from the analysis (we chose 50% inspired by the mixture modeling approaches used by FSL Melodic (Beckmann and Smith, 2004) to separate noise from signal, which uses 0.5 as a threshold to equally balance false-positives and false-negatives). This control excluded 3.0 ± 2.6% of voxels within the localizer mask for each subject, such that the final localizer masks contained 2907 ± 1129 voxels for each subject.

#### 2.5.4. Classifying voxels into veins and parenchyma

Voxels were classified as veins if the amplitude of measured oscillations was within the top 10% of all measured responses, subject to two constraints: (1.) the responses were in voxels in the pial surface, and (2.) the inverse tSNR of the voxel was below a threshold of 0.05 (Zhang et al., 2009), to avoid the accidental inclusion of bright voxels unlikely to be veins but that also displayed strong oscillations. This thresholding approach was visually inspected and compared against a high-resolution susceptibility-weighted image (SWI) acquired during pilot sessions in separate participants (data not shown). Since vein segmentation has been done based only on signal intensity in the past (Zhang et al., 2009), we also compared our approach to that prior method (Supplementary Figure 5). We note that this method does not allow us to identify intracortical veins; these results therefore focus on the effects of pial veins, which are expected to have the largest effects on BOLD response timing (Koopmans et al., 2010; Siero et al., 2009).

#### 2.5.5. Estimating the spatial spread of delays across V1

For each subject, the *x*, *y* and *z* coordinates of the V1 mask, in functional space, were mean centered and converted to millimeters. This yielded a position estimate from the center of V1, in millimeters, for each voxel within the mask. Since our imaging was performed such that slices were positioned parallel to the anterior-posterior axis of V1, we used the *y* -coordinate estimates as a proxy for the location along the anterior-posterior axis of V1.

#### 2.5.6. Statistical analysis

Statistical analysis was conducted using the R programming language (https://www.r-project.org) and the brms package (https://cloud.r-project.org/web/packages/brms). In particular, the brm function was used to fit Bayesian linear mixed models to the data to: (1.) estimate the effect of anatomical correlates on the observed amplitude, delay and temporal specificity of responses, and (2.) to estimate whether the amplitude and delay of responses at slow stimuli are related to observed amplitudes at fast stimuli. Weakly informative Gaussian priors were used by default and four Markov chains were run, each with 2000 iterations (discarding the first 1000 as warm-up), resulting in 4000 samples to approximate the posterior distribution of model parameters. The convergence of the Markov chains was assessed by computing the *R*^^^ statistic (Gelman-Rubin diagnostic) for each parameter. All chains for all models run in the current manuscript fully converged, as indicated by *R*^^^ statistics for all parameters almost identical to 1.

### 2.6. Simulating voxels with fast and slow frequency response

In order to simulate single-voxel frequency responses and compare them to experimental data, we simulated a range of HRFs modeled by a sum of two Gamma functions by sweeping across a range of peak delays (ranging from 3 to 6.5s) and full width at half maximum (FWHM) (ranging from 2.25 – 3.75s). For the second Gamma, which represents the undershoot, we kept the peak delay and FWHM fixed at 10s and 9.5s, respectively, with an amplitude of a fifth that of the positive peak. For the Siero HRFs, the amplitude of the undershoot was set to a tenth of the positive peak since that resembled better the figures shown in (Siero et al., 2011). For each combination of peak delay and FHWM, an HRF was sampled at 10-ms intervals, the result of which was then convolved with stimuli of different frequencies to generate a frequency response amplitude curve. We also simulated the frequency response of the canonical HRF (as described by (Glover, 1999)) and two HRFs that approximate the impulse responses for the pial and parenchymal visual cortex (as shown by (Siero et al., 2011)). These two HRFs had the following parameters for pial and parenchymal, respectively: time to peak = 3.57s and 3.30s, FWHM = 2.72s and 2.55s.

## 3. Results

### 3.1. Oscillatory stimuli enable measuring the temporal specificity of voxels in V1

To measure how well voxels in V1 can preserve high-frequency signals, we measured BOLD responses in the visual cortex induced by sinusoidally modulated visual stimuli at either slow or fast frequencies. This design allowed us to directly measure the ability of the vasculature to respond to increasingly faster stimuli, estimating the frequency response of each individual voxel. Oscillatory stimuli also have other advantages, such as minimizing neuronal nonlinearities at stimulus onset (Grill-Spector et al., 2006) and improving the separation between stimulus-evoked responses from background fluctuations (Kalatsky and Stryker, 2003). They are also simpler to analyze (Regan, 1966), since delays and amplitude can be unambiguously defined, whereas the onset and peak of transient responses, such as in blocked or event-related designs, are usually defined ad hoc and harder to compare across conditions. From the amplitude of oscillatory responses to different stimulus frequencies, we defined a “temporal specificity” metric, as a proxy of how much high frequency information is preserved in response to fast stimuli (Fig. 1).

**Figure 1:**
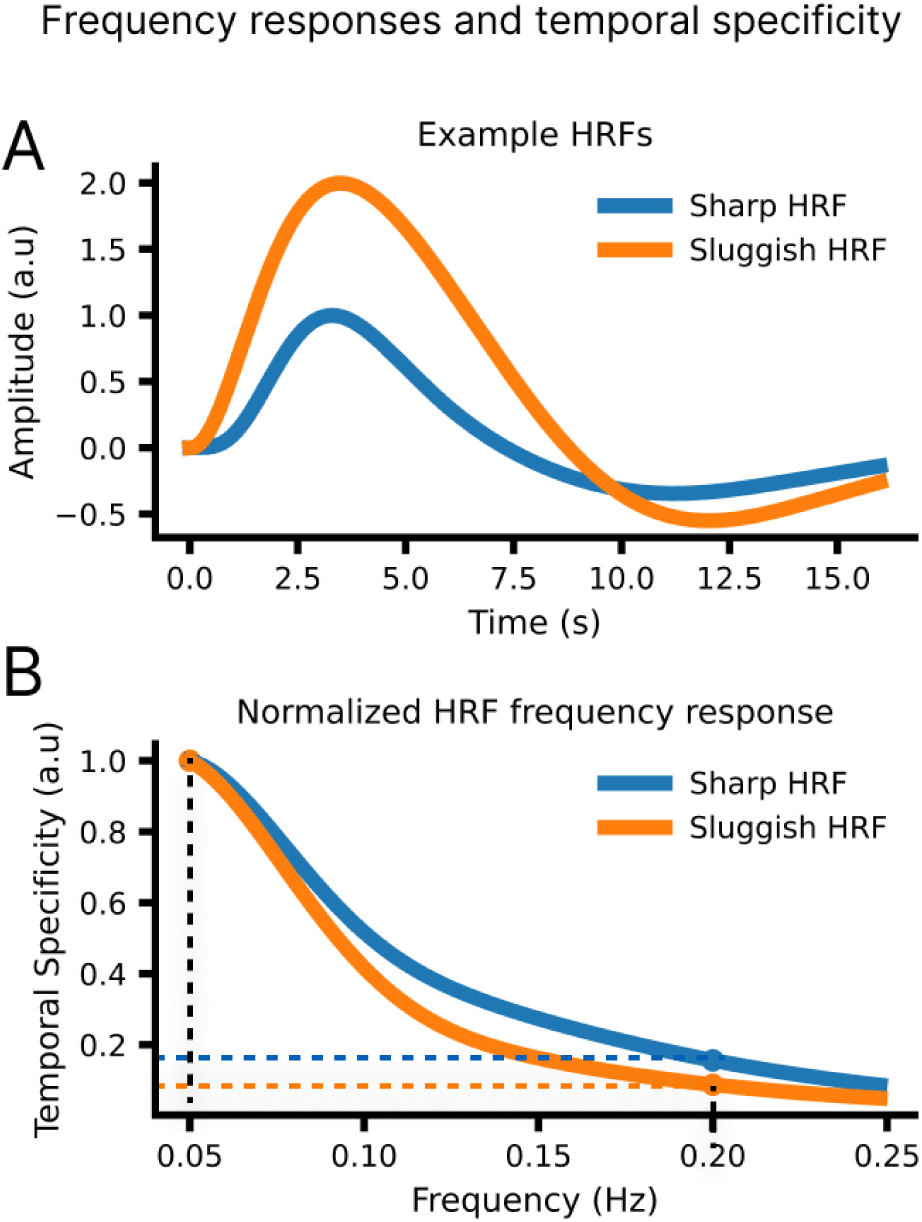
Temporal specificity is estimated by calculating the ratio of response amplitudes at different stimulus frequencies. A. A simulation of two example HRFs, one that is slow and has low amplitude (Sluggish HRF) and one that is fast and has a high amplitude (Sharp HRF). B. The frequency response is normalized to the reference response to 0.05Hz stimulation. The resulting frequency response can be used to compare which HRF produces relatively larger amplitudes at fast frequencies. This simulation illustrates that sharper HRFs should yield higher values of “temporal specificity”, i.e. with relatively high-amplitude responses at high frequencies.

### 3.2. High spatio-temporal resolution fMRI detects BOLD responses at the single-voxel level up to at least 0.20Hz

Since we aimed to investigate whether BOLD responses are linked to single-voxel vascular and anatomical properties, we first tested whether we could reliably identify BOLD fMRI responses to our oscillatory stimuli (Fig, 2, panel A) at the single-voxel level. We examined BOLD timeseries from all active voxels (Fig. 2, panel B), and the oscillatory patterns seen in individual voxels clearly demonstrated that responses were visible at the single-voxel level up to at least 0.20 Hz. From each voxel time-series, we calculated the mean cycle-locked response and extracted the amplitude and delay of the mean response for each voxel (Fig. 2, panel C). We observed a large heterogeneity in response properties across voxels, even at the single-subject level (Fig. 2, panel D), demonstrating the diversity of hemodynamic responses uncovered with high resolution imaging.

**Figure 2:**
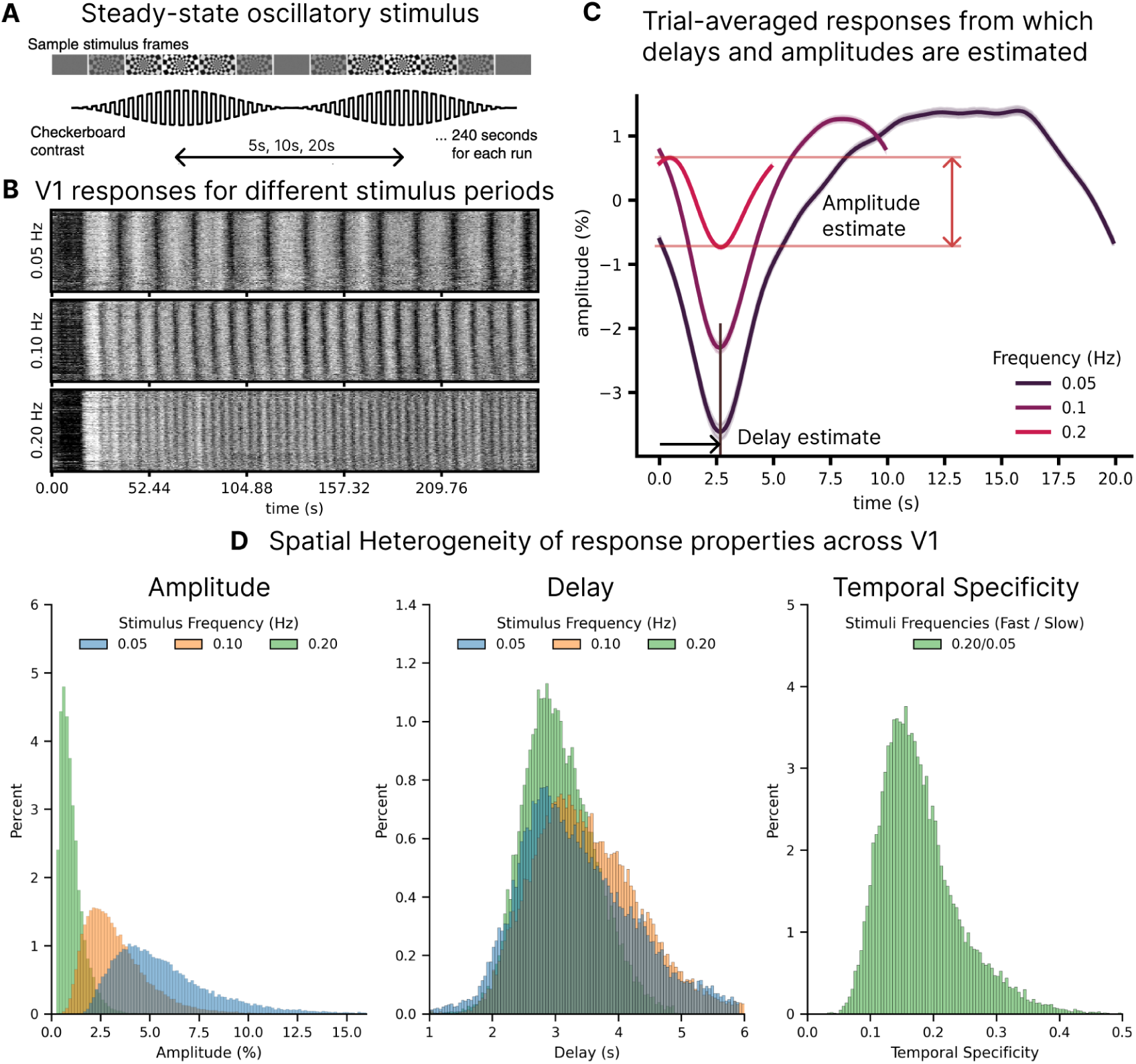
Single-voxel oscillatory responses can be reliably measured and show large heterogeneity. A. Continuous stimulus used to study the relationship between delays and hemodynamic variability: a flickering checkerboard with luminance contrast sinusoidally modulated with different frequencies. B. Carpet plots illustrating responses in the V1 ROI in an example subject, for three different frequencies. Each row represents a voxel within the localizer mask. C. Trial-averaged responses for the three stimuli in an example subject. Arrows indicate how delays and amplitudes are estimated. D. Histograms of the amplitude, delay and temporal specificity of responses show large heterogeneity even at the single-subject level.The y-axis show the percent of the total number of voxels analyzed.

### 3.3. Anatomical features influence the spatial patterns of delays and amplitudes and temporal specificity

Having extracted the oscillatory response in individual voxels, we next aimed to test how these responses were related to anatomical features within V1. We examined whether the estimated delay, amplitude and temporal specificity of single-voxel responses was linked to vascular-anatomical properties: cortical depth, vascular compartment (vein vs. parenchyma) and position along V1. We first inspected maps to qualitatively compare the spatial patterns of anatomy and vasculature against maps of delay and amplitude estimated for the response to different stimulus frequencies (Fig. 3, panel A). Measured patterns at the group-level aligned with previously reported BOLD properties (Koopmans et al., 2010; Siero et al., 2011, 2009): response amplitude and delay increased towards the pial surface, and responses were substantially larger in veins when compared to the parenchyma. We also observed a gradient of response properties across V1, with delay maps closely following the position along the anterior-posterior axis of V1. To quantify these spatial patterns, we conducted a Bayesian linear mixed effects analysis, testing whether anatomical predictors (cortical depth, position along the anterior-posterior axis of V1, and vascular compartment) significantly predicted fMRI response properties (delay, amplitude and temporal specificity in each voxel). We assumed that the absolute influence of cortical depth and position would vary for each subject (because of e.g. different cortical thickness or orientation of V1 with respect to B0 for each individual), thus modeling those as random effects. The model was fit against data from each frequency separately since the frequency effect is not expected to be linear.

**Figure 3:**
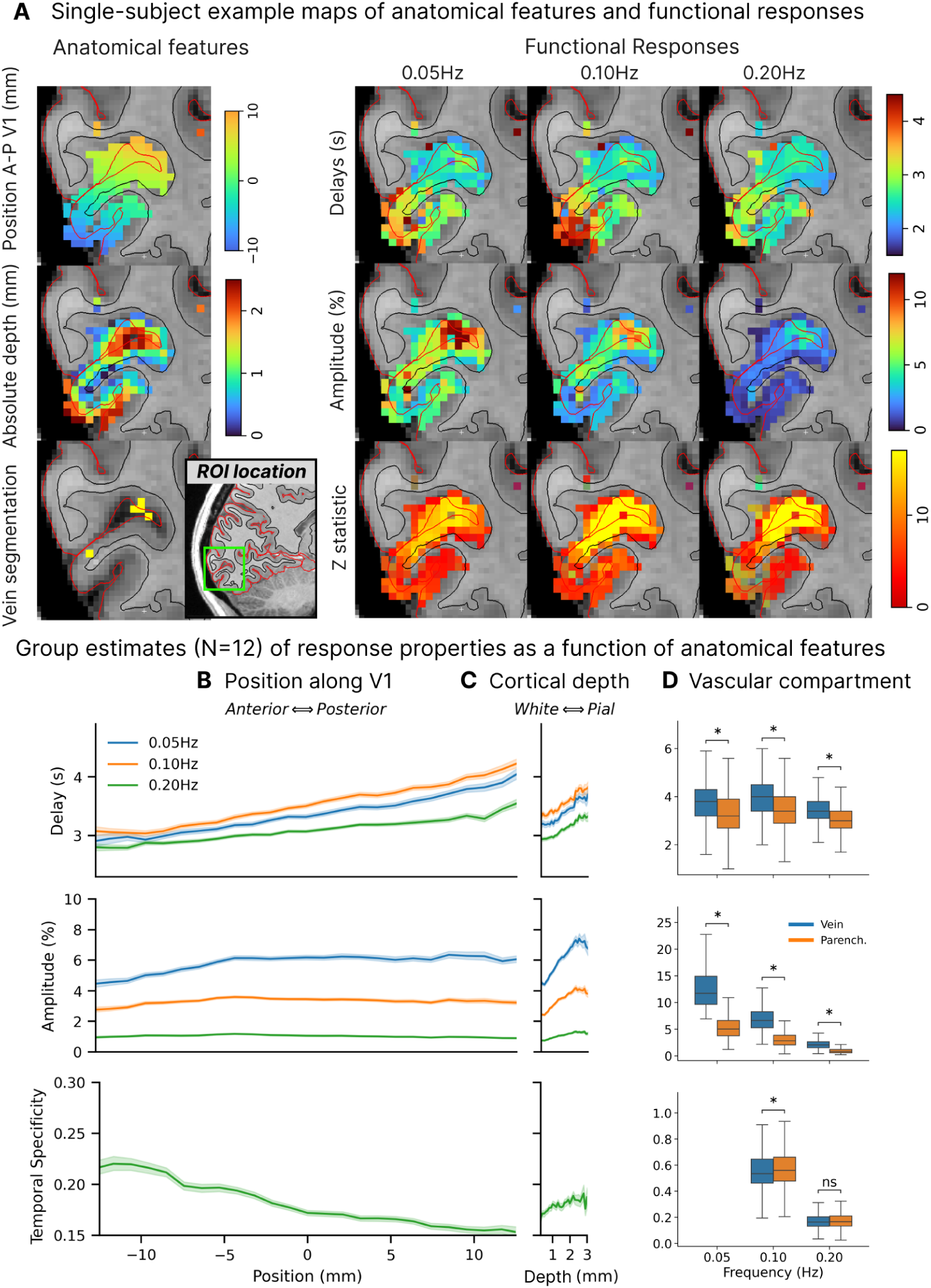
The amplitude, delay and temporal specificity of voxelwise responses are spatially structured. A. (left) Example maps of the three anatomical features used in this study: position along the A–P axis of V1, cortical depth and vein segmentation; Inset figure next to vein segmentation shows the ROI location in a sagittal slice on the right hemisphere over V1 for reference. (right) Maps of derived delay, amplitude and F/Z-statistic maps obtained across the three different stimulus frequencies. Maps are clearly spatially structured, with, i.e., the effect of depth and compartment clearly seen on amplitude maps, though the effect of position is less visually apparent at the single-subject level due to the smaller effect size and large subject variability (see Table 1 and Supplementary Figure 6). The effect of position, cortical depth and presence of veins can be qualitatively seen in the delay maps. B. Plots of delay, amplitude and temporal specificity (0.05Hz/0.20Hz) as a function of position along V1 (left) and cortical depth (right). Note that x-axis are on the same scale. C. Box plots of delay, amplitude and temporal specificity in veins and parenchyma for the three frequencies imaged. Stars indicate that differences were statistically significant (mixed-effects model, see Table 1). Panels B, C and D show data at the group-level. Shaded areas correspond to 95% CI over the mean over voxels.

**Table 1:** Mixed-effects model identifies strong effects of anatomical features on single-voxel fMRI responses. Weight estimates for delay, amplitude and specificity, for different stimulus frequencies as a function of cortical depth, position in V1, vascular compartment. Values in brackets represent 95% CI. (All temporal specificity values and gradients are multiplied by 100 for clarity.)

| ***Effect of anatomical correlates on response onset delay with respect to stimulus presentation*** |  |  |  |  |
| --- | --- | --- | --- | --- |
| Frequency | Delay at WM (ms) | WM → CSF (ms/mm) | Anterior → Posterior (ms/mm) | Impact of vein (ms) |
| 0.05 Hz | 3110 [2740, 3470] | 179 [123, 236] | 42 [28, 57] | 238 [206, 269] |
| 0.10 Hz | 3300 [2670, 3010] | 178 [127, 230] | 42 [33, 52] | 208 [181, 236] |
| 0.20 Hz | 2840 [2670, 3010] | 156 [103, 209] | 22 [15, 30] | 204 [182, 227] |

| ***Effect of anatomical correlates on signal amplitude*** |  |  |  |  |
| --- | --- | --- | --- | --- |
| Frequency | Amplitude at WM (%) | WM → CSF (%/mm) | Anterior → Posterior (%/mm) | Impact of vein (%) |
| 0.05 Hz | 4.38 [3.76, 5.00] | 0.77 [0.37, 1.18] | 0.04 [0.02, 0.07] | 6.99 [6.88, 7.10] |
| 0.10 Hz | 2.42 [2.00, 2.83] | 0.45 [0.27, 0.61] | 0.00 [-0.01, 0.01] | 3.74 [3.68, 3.81] |
| 0.20 Hz | 0.74 [0.62, 0.86] | 0.14 [0.07, 0.22] | -0.007 [-0.012, -0.001] | 1.19 [1.16, 1.21] |

| ***Effect of anatomical correlates on temporal specificity (multiplied by 100)*** |  |  |  |  |
| --- | --- | --- | --- | --- |
| Frequency | 100 × specificity at WM | WM → CSF (mm<sup>-1</sup>) | Anterior → Posterior (mm<sup>-1</sup>) | Impact of vein |
| 0.10 Hz | 55.07 [50.96, 59.33] | 1.04 [0.05, 2.05] | -0.44 [-0.27, -0.62] | -1.06 [-1.61, -0.50] |
| 0.20 Hz | 16.88 [15.29, 18.52] | 0.39 [-0.06, 0.86] | -0.25 [-0.34, -0.17] | -0.12 [-0.34, 0.11] |

#### 3.3.1. Effect of anatomical covariates on hemodynamic delays

Using these model estimates, we first examined how each anatomical feature related to the delay of the fMRI responses. The effect of cortical depth on delay was similar for all frequencies, with delay increasing from the white to the pial surface by 170 ms/mm on average. The presence of a vein was associated with an additional 215 ms delay, beyond the effect of cortical depth. The magnitude of the effect of vascular compartment on delay was also consistent across different stimulus frequencies (Table 1). The anatomical position was also a significant predictor, with delays increasing towards posterior V1 (from the calcarine sulcus towards the occipital pole), however, this relationship showed a dependence on stimulus frequency, with weaker effects at the faster frequencies (from 42 ms/mm at 0.05 and 0.10 Hz to 22 ms/mm at 0.20 Hz, Table 1). These results demonstrated that the depth, vascular anatomy, and position of voxels each contributed to its response delay.

#### 3.3.2. Effect of anatomical covariates on response amplitude

We next investigated the spatial pattern of response amplitudes, investigating which voxels had the largest responses. Amplitudes dropped substantially when using faster stimuli, as expected from prior work (Lewis et al., 2016): the mean amplitude at the white matter at the center of V1 was: 4.38% (95% CI: 3.76 to 5.00%) for the 0.05Hz stimulus, 2.42% (95% CI: 2.00 to 2.83%) for the 0.10Hz stimulus, and 0.74% (95% CI: 0.62 to 0.86%) for the 0.20Hz stimulus. The response amplitudes were highly variable, and some voxels showed extremely large responses; notably, individual single-voxel responses reached as high as 60.84% for one participant, with the largest voxelwise amplitudes averaging 33.56% ± 16.25% across all 12 participants. Response amplitudes increased substantially as a function of cortical depth and were heavily influenced by the presence of a vein with substantial amplitude increases (Table 1). This is consistent with prior studies observing the largest responses at the cortical surface and veins (Siero et al., 2011, 2009). Importantly, amplitudes were strongest in veins for all stimulus conditions (Figure 3, panel D).

While response amplitudes were consistently related to cortical depth and compartment, the effect of position differed qualitatively across stimulus frequencies. For the 0.05Hz stimulus, the amplitude significantly increased towards the posterior part of V1 by 0.04%/mm (95% CI: 0.02 to 0.07%) being largest closer to the occipital pole. For the 0.10Hz stimulus, no significant trend was observed in the amplitude as a function of position (0.00%, 95% CI: −0.01 to 0.01%). For the 0.20Hz stimulus, the amplitude significantly decreased towards the posterior part of V1 by 0.007% (95% CI: 0.001 to 0.012%) being largest deep into the calcarine sulcus (Figure 3, panels B and C). This pattern suggests that regions with fast responses may be spatially structured in such a way as to yield distinct spatial maps under different stimulus frequencies, as the slow stimulus biases responses to be stronger in posterior V1 whereas fast stimulus biases them to be larger in anterior V1. Nonetheless, we note that the effect of spatial location on response amplitude, albeit significant, is much smaller in magnitude than the effect of cortical depth (20-fold larger) or presence of a vein (174-fold larger), hence not qualitatively visible on an individual amplitude map.

#### 3.3.3. Effect of anatomical covariates on temporal specificity

Finally, to test our hypothesis that the temporal precision of functional MRI is also spatially structured, we investigated the spatial structure of temporal specificity. A temporal specificity value of 0 indicates a zero amplitude at the fast frequency, and a value of 1 signifies that the response amplitude remains constant across slow and fast frequencies. Higher values imply voxels with relatively stronger responses to the faster stimulus, indicating that they more effectively retain temporal information and are thus more “temporally specific”.

We found no significant effect of cortical depth on temporal specificity, with a change of +0.0039 mm^-1^ (95% CI: −0.0006 to 0.0086), a small number that suggests voxels sampling the pial surface preserves only about 0.4% more amplitude at 0.20Hz compared to 0.05Hz than those that sample the white matter. We point out, however, that the magnitude of the effect of cortical depth should be interpreted with care. We found no statistically significant linear effect of cortical depth, but cortical depth could have a more nuanced effect on specificity. The trend in these data suggested that temporal specificity may increase slightly towards the pial surface until depths of about 1 mm (our imaging resolution), an effect likely related to partial volume effects, but then flatten out (Fig. 3, panel B). This suggests that the effect of cortical depth on specificity is not strictly linear, and also that comparing temporal specificity differences between “deep” and “superficial” bins of the cortical ribbon may lead to statistically significant results based on where the threshold is placed that defines whether a voxel is in the deep or superficial bin.

We next examined the effects of veins, and found that voxels with veins had moderately lower temporal specificity in the 0.10Hz stimulus (reduction of −0.0106 (95%CI: −0.0161 to −0.0050)), and a similar trend was seen in the response to the 0.20Hz stimulus (reduction of −0.0012 (95%CI: −0.0034 to 0.0011)). These results demonstrated a subtle effect of veins on temporal specificity, with slightly decreased temporal specificity in the veins.

In contrast to the subtle effects of depth and vascular anatomy, we observed a strong relationship between anatomical location and temporal precision, with a steep reduction in specificity from anterior to posterior V1 by 0.0025 mm^-1^ (95%CI: 0.0017 to 0.0034). To exemplify the size of this effect we can consider two arbitrary voxels, a voxel A in anterior V1 about 2cm away along the A–P axis of V1 from a voxel B in posterior V1. If they show the same response amplitude of 10% at 0.05Hz stimulation, our analysis suggests that at 0.20 Hz, voxel A would have an amplitude of about 2%, and voxel B an amplitude of 1.5%, a large difference due to positional effects alone. Our analysis also revealed significant between-subject variability (0.0283, 95%CI: 0.0192 to 0.0425, values larger than the estimated effect of depth or compartment) in mean temporal specificity, which suggests that fast fMRI responses may be more prominent in some but not other individuals, perhaps due to variability in properties of blood and vasculature or perhaps other aspects of baseline physiology or neuronal activity.

### 3.4. Functional responses to slow stimuli are linked to temporal specificity

These results demonstrated that there is a strong relationship between vascular-anatomical properties and BOLD amplitude and delay estimates, and a more complex relationship to temporal specificity, which is substantially influenced by anatomical location along the A–P axis of V1. Nonetheless, in practice not all experiments can be conducted at a sufficiently high resolution to allow for cortical depth analysis or vascular segmentation, nor does every experimental session provide sufficient time to acquire multiple runs to map out temporal specificity. Furthermore, the fact that there is strong subject variability suggests that using anatomical covariates alone is not sufficient to fully explain responses at the single-subject level. Therefore, we also investigated whether the responses to a fast stimulus were related to the responses to the slower stimulus, since slow task designs are ubiquitous in fMRI experiments, and would enable subject-specific analyses to account for local temporal specificity potentially also outside of V1. To do so, we fit a mixed-effects linear model to test whether the amplitude at the 0.20Hz stimulus was linked to the amplitude and delay from the 0.05Hz stimulus (which are simple to obtain in a short experiment).

This analysis confirmed a significant positive relationship between the amplitude at the slow stimulus (0.05Hz) and the fast stimulus (0.20Hz): the group-level estimate for the effect of amplitude was 0.24 (95% CI: 0.22 to 0.27), meaning that response amplitudes at 0.20Hz stimulus will range between approximately 22–27% of the response amplitude at 0.05 Hz. The effect of delay alone was not significant (0.01; 95% CI: -0.03 to 0.06), meaning that knowledge of the delay alone is not sufficient to predict the amplitude at fast frequency, as expected since baseline amplitudes are variable across voxels. Importantly, however, there was a strong negative interaction between the delay and amplitude, of −0.020 (95% CI: −0.022 to −0.019). This negative interaction implies that given knowledge of a voxel’s response amplitude at 0.05 Hz, short delays will predict higher relative amplitudes at 0.20 Hz. In other words, for two voxels with identical response amplitudes at 0.05 Hz, we expect a larger amplitude at 0.20 Hz for the voxel with shorter delay, meaning that delay is correlated with temporal specificity.. This is illustrated in the scatter plot of figure 4 (panel A), which shows, for each voxel, amplitudes at 0.05 Hz and 0.20 Hz color-coded by their delay at 0.05 Hz. We found that temporal specificity, (the ratio of the responses to the 0.20Hz and 0.05Hz stimuli) is related to the slope, and datapoints with different delay values appear to lie along straight lines with different slopes, with the shorter delay values exhibiting a steeper slope (higher temporal specificity) and longer delay values exhibiting a shallower slope (lower temporal specificity), indicating an anti-correlation between delay and temporal specificity. This highlights that short delays associate with higher temporal specificity, and thus better preservation of fast responses (see also Supplementary Figure 7). The effect is also shown qualitatively in two individual subject maps (figure 4 B) with delay maps showing similar spatial patterns to those of temporal specificity. This holds even though these two example subjects have clearly different anatomy. These results demonstrate that, in addition to our observation that anatomical covariates can explain substantial variance in temporal specificity, the properties of the functional response to slow stimuli are also strongly related to temporal specificity, which in future studies could provide a complementary approach to identify fast voxels in addition to anatomical criteria.

**Figure 4:**
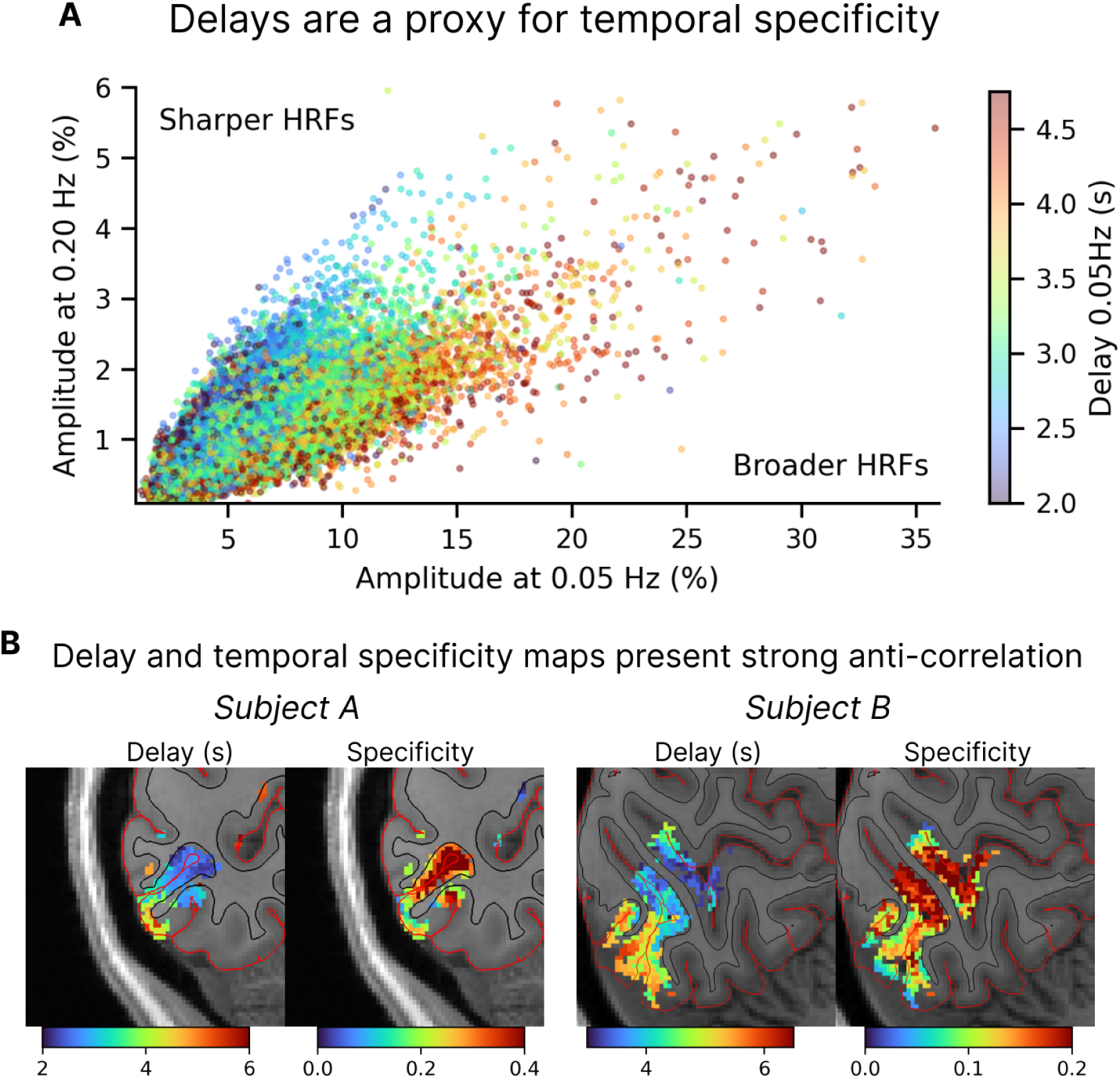
Relationship between delays and temporal specificity. A. The amplitude values at two different stimuli color-coded by the delay (a measure of hemodynamic latency). The delay and amplitude of the response to a slow stimulus are thus linked to the amplitude of the response to a fast stimulus (n=12 subjects). B. Maps of the hemodynamic delays measured with the 0.10Hz stimulus and temporal specificity estimated as the ratio of amplitudes of the responses to fast and slow stimuli (0.20 Hz / 0.05 Hz), shown for two example subjects with distinct cortical folding. A clear anti-correlation can be seen where voxels with the lowest delays tend to also have the highest specificities.

### 3.5. Temporal precision may be enhanced by hemodynamic non-linearity

Overall, our results demonstrated relatively robust responses to the fastest 0.20 Hz stimulus, indicating that voxels that respond fast are widespread in V1. The robust, fast responses could reflect HRFs that are generally sharper and more precise than expected, or could reflect voxels with nonlinear hemodynamics, such that faster stimuli elicit more temporally precise HRFs. We therefore investigated whether a single HRF (i.e., a linear model) could be used to predict the amplitude behavior observed in our experimental data. HRFs become faster and sharper as the stimuli become shorter (Lewis et al., 2018; Yeşilyurt et al., 2008), so we hypothesized that responses to rapidly oscillating stimuli would in part reflect a nonlinear change in voxelwise HRFs. To answer this question we compared the temporal specificity at 0.20 Hz and at 0.10 Hz against a family of physiologically plausible HRFs parameterized by their full-width at half-maximum and their peak delay, the canonical HRF (Glover, 1999) and two compartment-specific HRFs, parenchymal and venous (Siero et al., 2011). As hypothesized, we found that no single HRF was capable of satisfactorily predicting the amplitude across frequencies observed in our data, as the measured responses to the high-frequency stimulus were larger than models predicted. Instead, we found an apparent “acceleration” of HRF as frequency increases (Figure 5): slower HRFs with broader FWHMs (Fig. 5, in blue) better predicted the responses to the 0.10Hz stimulus, and faster HRFs with narrower FWHMs (in red) better predicted the responses to the 0.20Hz stimulus. In particular, the canonical HRF clearly failed to predict responses to either stimulus, and the Siero HRFs best predicted responses to 0.20Hz. These results suggest that a nonlinearity manifesting as an acceleration of the HRF with faster stimulus frequencies may have contributed in part to the fast responses observed in our study.

**Figure 5:**
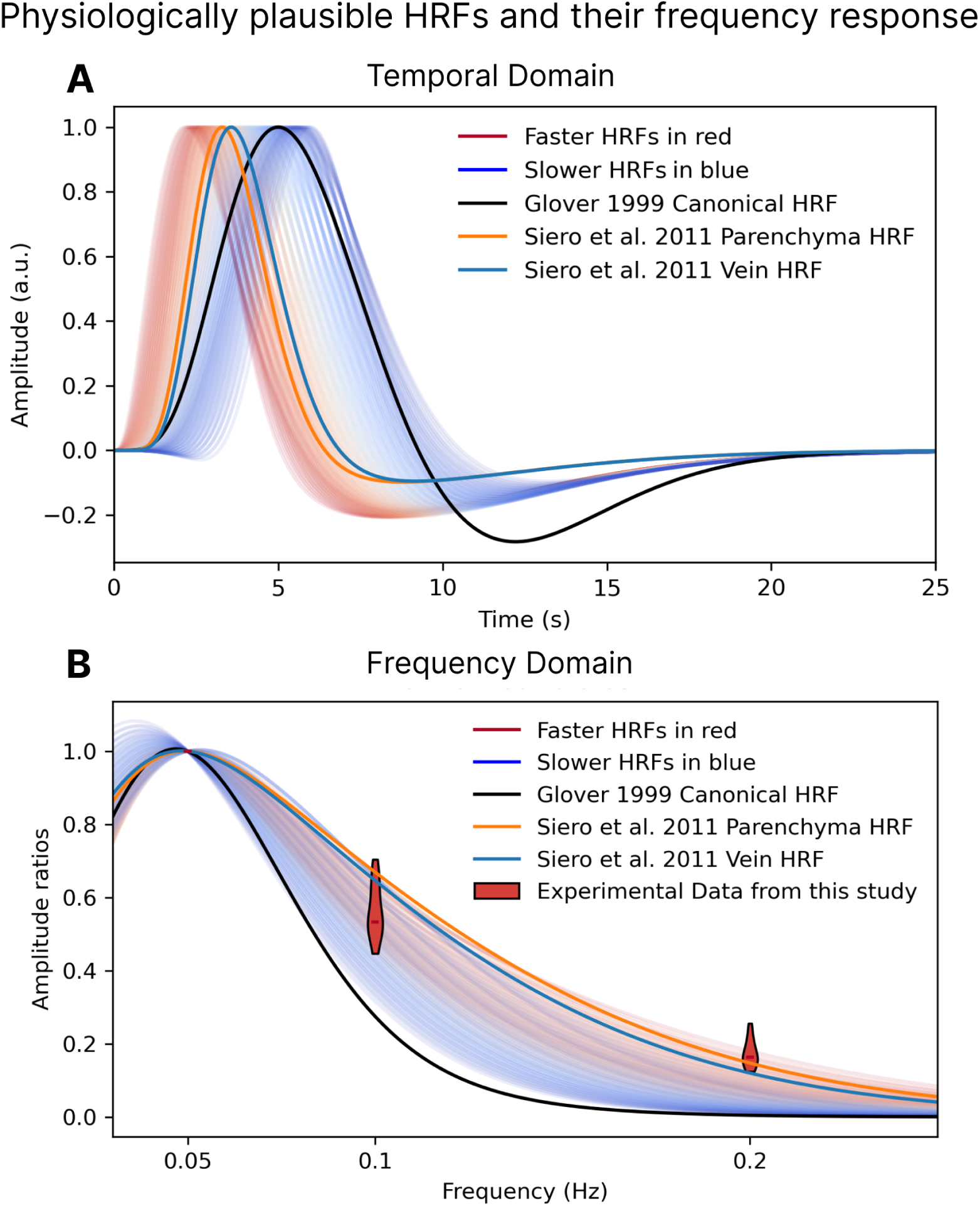
Different HRFs predict the fMRI responses to 0.10Hz and 0.20Hz stimuli. A. Physiologically plausible HRFs generated by varying the FWHM (2.25–3.75 s) and onset delay (2.3–5.4 s) of a double-Gamma function. Values normalized such that maximum positive amplitude is equal to 1. B. Curves of normalized frequency responses for the family of HRFs shown in panel A. No combination of FWHM or onset delay generates a frequency response that matches the observed amplitude ratios. Faster HRFs are shown in red, slower in blue, with the gradient of colors changing from darkest blue to darkest red as the HRFs become faster. The red violin plots show the temporal specificity estimates for each frequency. The dark line represents the canonical HRF, which clearly underestimates response amplitudes for both the 0.10- and 0.20Hz stimuli. In orange and blue are the Siero HRFs which best predict the 0.20Hz responses but overestimate the amplitude of 0.10Hz responses.

## 4. Discussion

The interpretation of fMRI studies is critically influenced by hemodynamic response properties: if voxels with different response properties are spatially organized in distinct regions of the cortex, this can be reflected in statistical activation maps. Here we showed how the properties of V1 responses are influenced by anatomical features. Cortical depth, veins, and anatomical position all influenced single-voxel response properties. Cortical depth and the presence of veins strongly influenced delays and amplitudes, but only weakly influenced temporal specificity. Surprisingly, we found that the strongest predictor of temporal specificity in V1 was anatomical position across the A–P axis. This spatial gradient strongly dominated temporal specificity and produced distinct spatial patterns of response amplitudes when using fast, as opposed to slow, stimuli.

### 4.1. Anatomical features associated with single-voxel responses

In accordance with previous literature, we found that hemodynamic delays measured with BOLD within the cortical gray matter increase from the white matter to the pial surface (Havlicek and Uludağ, 2020; Markuerkiaga et al., 2016; de Zwart et al., 2005), and increase in the presence of veins. Response amplitudes also increased from the white matter to the pial surface and increased substantially in the presence of veins (Koopmans et al., 2010; Siero et al., 2011, 2009). It is worth noting, however, that the amplitude of parenchymal voxels—both those intersecting the white matter and those intersecting the pial surface—can be strikingly similar when compared to amplitude values found in veins. This suggests that, despite a correlation between cortical depth and vascular compartment, not all voxels at the pial surface should be expected to have a large BOLD amplitude—only those in the proximity of large pial veins. The effects of pial veins could be relatively sparse when imaging at high spatial resolution, because not all voxels at the pial surface will contain large draining veins due to the sparsity of these large veins on the surface of the brain.

Our analysis also measured the temporal specificity of fMRI responses, testing how robustly each voxel could retain temporal information at faster frequencies. Our results demonstrated that spatial position within V1 has a large effect on temporal specificity, and far outweighs the effects of veins or cortical depth, with voxels in anterior V1 being relatively more sensitive to fast stimuli. Even though positional effects had been observed in the context of the entire human cortex (Handwerker et al., 2004), we find local hemodynamic differences to be substantial even within distances as small as 2–3 cm within the primary visual cortex. We tested whether an interaction between veins and location could explain this effect, but found this not to be the case. Positional effects were also preserved when excluding veins from the analysis, suggesting that large veins alone do not explain this gradient (Supplementary Figures 8 and 9).

The local hemodynamic variability suggests that biases could emerge when the intrinsic temporal properties of voxels align or misalign with the stimulus frequency or experimental design. While designs that use high frequencies or approaches like event-related and naturalistic designs may be more biased towards fast voxels, so are designs that use low frequencies or long duration blocked stimuli likely to be biased towards slower voxels. Therefore, careful consideration must be given to positional effects, as they could lead to discrepancies in amplitude estimates across regions that are unrelated to the neuronal effects of interest. Our results focus on V1 and identify a spatial gradient of temporal specificity within this region. A key next step will be to investigate whether analogous gradients are present in other cortical regions. One potential approach our results suggest is to use delay maps as a proxy for temporal specificity.

### 4.2. On the striking and widespread heterogeneity of responses within V1

The spatial heterogeneity of temporal properties of BOLD responses across V1 was striking. Indeed, earlier studies had already exhibited a structured spatial patterns of BOLD responses across the visual cortex, suggesting these effects may be broadly present in other datasets. For example, analysis of resting-state and task-driven fMRI datasets have identified multiple visual networks in the human brain (Smith et al., 2009; Yeo et al., 2011), namely a medial, an occipital and a lateral network. Since functional connectivity estimates are delay-sensitive, it could be that the exact boundaries at which networks segregate are influenced by differences in their hemodynamic response properties even if the underlying neuronal activity were synchronous. Although prior studies have not specifically tested this effect, their results often display a spatial pattern that resembles our observed delays (Amemiya et al., 2020; Bailes et al., 2023; Lewis et al., 2016, fig. 7). These experimental observations all converge on the notion that local hemodynamic differences are widespread, particularly when imaging at high resolution. A distinctive aspect of our work that allowed us to study these effects was the use of high temporal and spatial resolution imaging, and an analysis pipeline focused on the single subject, extracting data in each subject’s native space to minimize loss of specificity due to smoothing and interpolation (Wang et al., 2022).

### 4.3. Potential causes for the heterogeneity in specificity and its relationship to response delays

What mechanisms could underlie this heterogeneity in temporal specificity, and why is it that differences in delays across cortical depths and vascular compartments are pronounced but these anatomical features only weakly relate to temporal specificity? Our data are not able to address this, but some possibilities can be considered.

Specifically, the dominant mechanism contributing to delays may differ when considering either position within a cortical region or cortical depth at a given position. While both factors contribute to the observed delay, it is helpful to consider them separately. The delay at a given position is determined by the pial macrovascular anatomy and physiology, whereas the delay at a given depth is determined by the intracortical microvasculature. These delays at a given position therefore will depend on arterial arrival times and venous drainage times, which can vary substantially across the cortex. While the BOLD signal mainly reflects venous effects, the timing of the hemodynamics is also affected by arterial response timing. Although arterial arrival times — which do follow the A–P axis as blood is delivered to the visual cortex via the posterior cerebral artery — may partly explain our findings, functional hyperemia is known to propagate upstream from the sites of activation, thus BOLD delays should not unambiguously track blood delivery times. Delays could also reflect venous effects towards the inferior and superior sagittal sinuses, which also run in the A–P direction. Delays across cortical depths, however, are expected to be consistent throughout V1 because the microvascular architecture is similar across V1 (Weber et al., 2008). Here, arterial timing is also relevant for BOLD response timing, and while there may be some anatomical asymmetries in the diving arterioles and ascending venules (Duvernoy et al., 1981) that may lead to local asymmetries in the timing of blood supply and drainage across cortical depths, it is unknown whether such asymmetries in angioarchitecture vary systematically within V1. Overall, the distinct mechanisms underlying delays varying across position versus delays varying across cortical depths may each exert some effect on temporal specificity. Future work could investigate these macrovascular and microvascular components to identify mechanisms underlying the heterogeneity of the observed temporal properties seen in our data.

Finally, we believe the evidence suggests that hemodynamic properties are more likely to be the primary driver of our observations; we cannot fully rule out some contribution from neuronal effects, but prior studies using direct neuronal measurements suggest that neuronal timing should not differ enough across V1 to cause delay differences as large as the ones we observe (>1 s). Evoked potentials measured in foveal, parafoveal, and peripheral representations within visual cortex show similar timing and amplitude, with timing differences between foveal and peripheral representations being smaller than 50 ms (Hansen et al., 2016). In addition, our stimulus excluded both the fovea and far periphery of the visual field, reducing neuronal variance in the responses across the stimulated region of V1. However, the amplitude of the neuronal activity could vary across V1, for example due to different sensitivity to luminance, imperfect scaling of our radial checkerboards, or slight differences in the spatial frequency across regions, which could influence nonlinearity of the fMRI response and alter spatial patterns. Given the large timing delay we report across regions, of over a second, it is unlikely that neural differences could fully explain these results, but neuronal effects may nevertheless have some influence on the observed amplitudes and thus the spatial patterns of temporal specificity.

### 4.4. Implications for laminar fMRI

The results we presented regarding the impact of cortical depth and vascular compartment on responses agree with many previous reports from the literature (Gati et al., 1997; Lai et al., 1993; Siero et al., 2011, 2009). They corroborate the importance of accounting for venous effects and depth-dependent biases on the BOLD signal. Nonetheless, despite potentially biasing analysis, since the amplitude of veins can be almost an order of magnitude larger than those of parenchymal voxels, the venous signal could still be exploited as a sentinel for neuronal activity at high frequencies, even if their spatial location is less specific. Veins are slower, but their larger amplitude may balance out their lower temporal specificity, and as such, at least up until frequencies of 0.20Hz, and potentially even higher, the largest amplitudes should still be expected in venous voxels.

### 4.5. Implications for fast fMRI

While the heterogeneity of single-voxel responses can pose analytic challenges, it also reflects an opportunity for fast fMRI: fast hemodynamics are widespread and common in the cortex. Identifying the location of fast voxels could therefore be highly advantageous for conducting successful fast fMRI experiments. Due to the small amplitude of fast responses, an ideal imaging approach would focus on areas where responses are fast enough to be visible or where the baseline BOLD amplitude is overwhelmingly large, such as in individual venous voxels. Our results demonstrate how anatomy and delayed functional responses are linked to temporal specificity and could guide localizer and ROI selection to enhance detectability of fast responses. Taking advantage of the location of fast voxels may require revising strategies for data preprocessing and analysis, possibly requiring more detailed anatomical imaging to create finer masks and regions-of-interest (ROIs) and data registration and averaging techniques informed by anatomical and vascular segmentation, which are currently being developed in the context of high spatial resolution imaging as well (Blazejewska et al., 2019; Wang et al., 2022). Moreover, when modeling fast fMRI using linear models adopting a single HRF, our data supports employing faster hemodynamic response functions HRFs for capturing the dynamics, such as those measured by Siero et al. 2011. Those could be used as updated “canonical-fast” response functions for high-resolution fast fMRI, since the conventional canonical HRFs clearly underestimate the amplitude of responses to faster frequencies. Finally, another implication for fast fMRI studies is that as the frequency of stimulation increases, the relative impact of veins becomes smaller. This result suggests that improved spatial specificity achieved through reduced contribution from draining veins can sometimes accompany improved temporal specificity in fast fMRI.

### 4.6. Identifying mechanisms of fast responses and avenues for further study

In future studies, it would be crucial to explore mechanisms contributing to the fast and slow responses observed here in the visual cortex. One possible avenue of investigation includes the use of fast functional Magnetic Resonance Angiography (fMRA) to image blood velocity, flow and potentially vessel dilation (Bizeau et al., 2017; Cho et al., 2012, 2008) to understand the underlying physiology. Furthermore, examining perfusion or blood volume changes could provide insights into how the various hemodynamic components interact to produce the fast BOLD responses. Expanding the application of fast fMRI to other brain regions, as proposed by Hodono et al. 2022, may help elucidate the generalizability of these findings. Additionally, further investigations on the impact of baseline blood flow, as in combining gas challenges with task fMRI, could shed light on the interaction between baseline brain activity and the speed of the hemodynamic responses. For instance, it would be worthwhile to test whether using fast stimuli with hypocapnia (which has been shown to accelerate BOLD responses (Cohen et al., 2002)) could lead to even stronger responses to fast stimuli. Since our results were obtained in healthy young adults, future studies should also examine whether these effects generalize to clinical or aging populations, particularly since vascular anatomy and physiology can change substantially with age (D’Esposito et al., 2003). Lastly, it is worth noting that recent advances in MR instrumentation and processing methods (Bates et al., 2023; Feinberg et al., 2023; Vizioli et al., 2021) could soon allow the direct imaging of even faster frequencies. These advances in imaging, when combined with advances in modeling of fast responses (Polimeni and Lewis, 2021), will further extend the capabilities of fMRI towards its biological limits.

## 5. Data and Code Availability Statement

Datasets used for this study will be made available upon publication in anonymized and deidentified form. Analysis code will be made available under https://github.com/dangom/gomez2024temporal. Code for the visual stimulation software can be found under In case of issues or data related queries please contact the corresponding author.

## 6. Author Contributions

**Conceptualization** DG, JP, LL

**Methodology** DG, JP, LL

**Software** DG, JP

**Validation** DG, LL

**Formal Analysis** DG, JP, LL

**Investigation** DG, JP, LL

**Resources** JP, LL

**Data Curation** DG

**Writing - Original Draft** DG, LL

**Writing - Review and Editing** DG, JP, LL

**Visualization** DG

**Supervision** JP, LL

**Project Administration** DG, LL

**Funding Acquisition** JP, LL

## Acknowledgements

We would like to thank Nina Fultz and Kyle Droppa for support during data acquisition. We would like to thank Grant Hartung and Jörg Pfannmöller for useful discussions during the preparation of this manuscript.

