## Supplementary Materials for "The temporal specificity of BOLD fMRI is systematically related to anatomical and vascular features of the human brain"

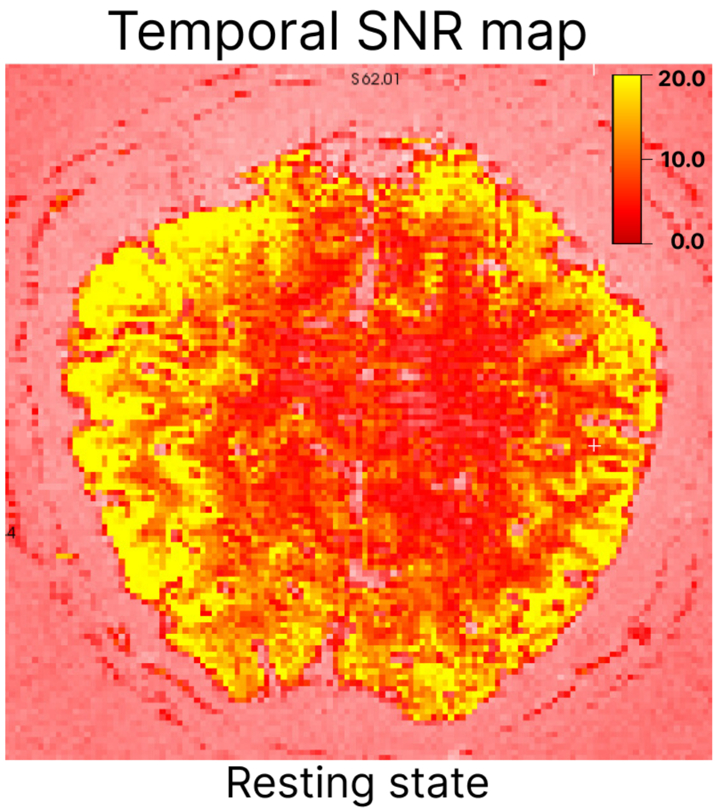


***Supplementary Figure 1. Temporal SNR for data quality assessment.*** Temporal SNR calculated as the temporal mean divided by the temporal standard deviation of the time-series data. Shown is a single coronal slice positioned over the calcarine sulcus. Colorbar indicates tSNR values ranging from 0–20. The tSNR was calculated over a resting-state run of same duration as the task run (257 s), acquired in one participant. The average tSNR within the ROI was 13.

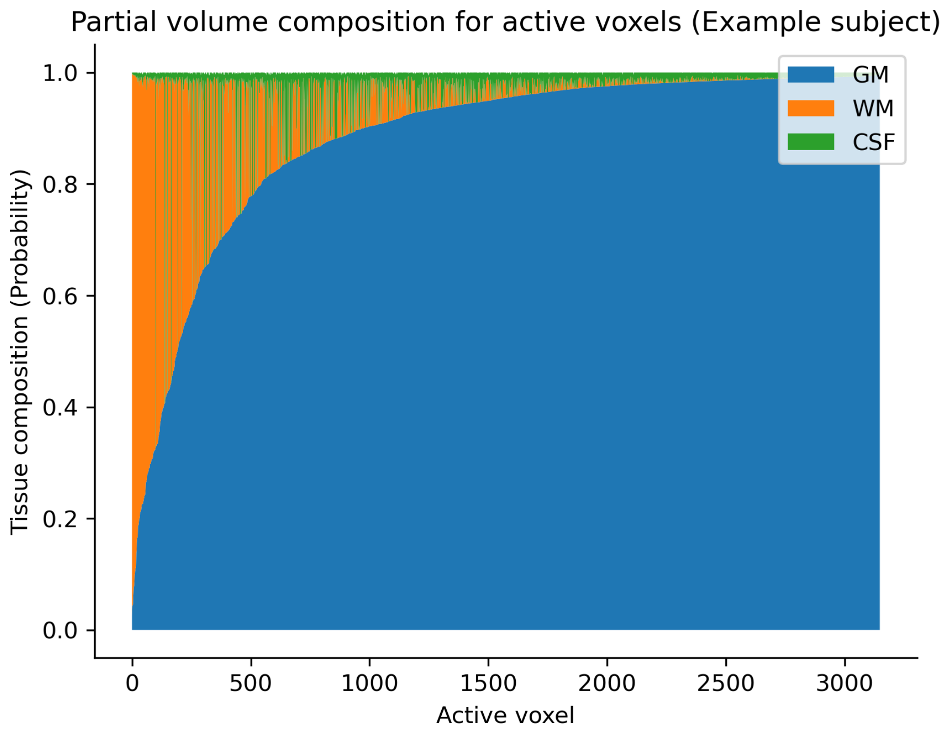


***Supplementary Figure 2. Partial volume analysis showing the tissue composition of active voxels of an example subject.*** The majority of voxels contain mostly gray matter (shown in blue), with remaining voxels containing increasingly larger WM and CSF contamination. These partial volume calculations were obtained by using SPM cortical segmentation projected onto the functional data's native space.-We found that 85.7% of active voxels (across all subjects) contain more than 50% gray matter, with the remaining voxels containing increasingly larger contributions of WM and CSF.

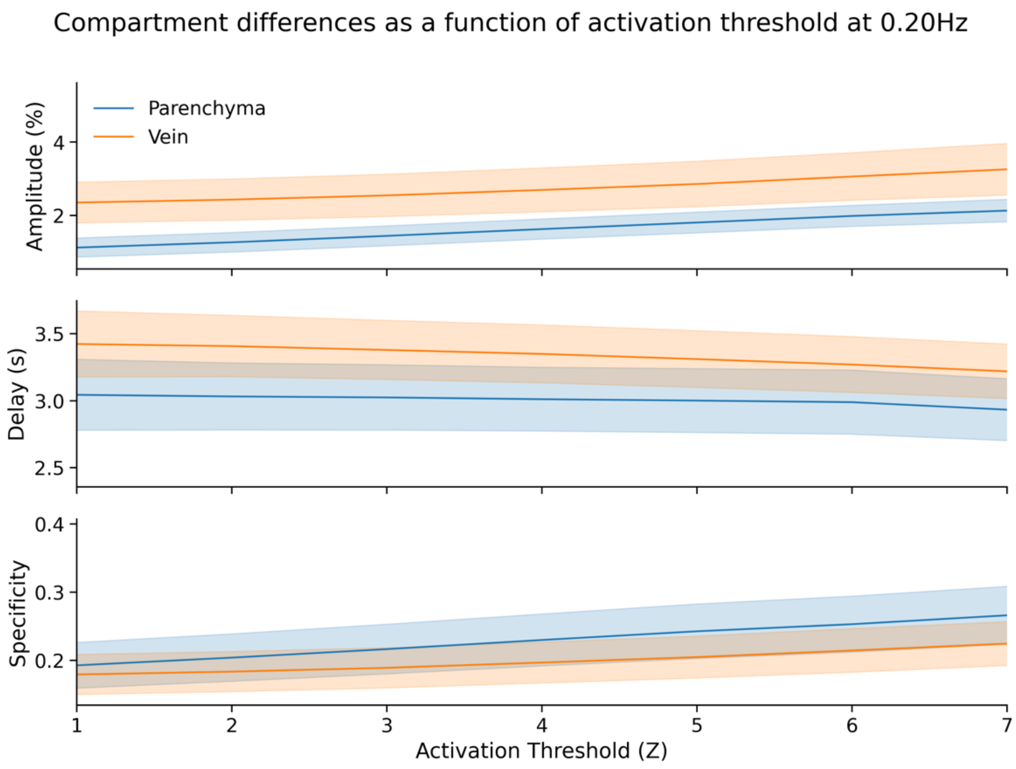


***Supplementary Figure 3. The difference between parenchyma and veins is qualitatively similar across voxel selection thresholds.*** Plot shows amplitudes, delays and temporal specificity as a function of activation threshold used in the localizer to select voxels for further analysis. Values are taken for the 0.20-Hz stimulus, the most sensitive to thresholding since amplitudes are the smallest amongst each stimulus condition. Changes in activation threshold influence the absolute values of mean amplitude, delay and specificity, but do not cause the direction of effects to change, i.e., veins always have larger amplitudes. Increasing the z-threshold biases estimates to faster voxels with shorter delays and larger amplitudes, while removing slower voxels with smaller amplitudes. Shaded areas represent standard deviation across voxels.

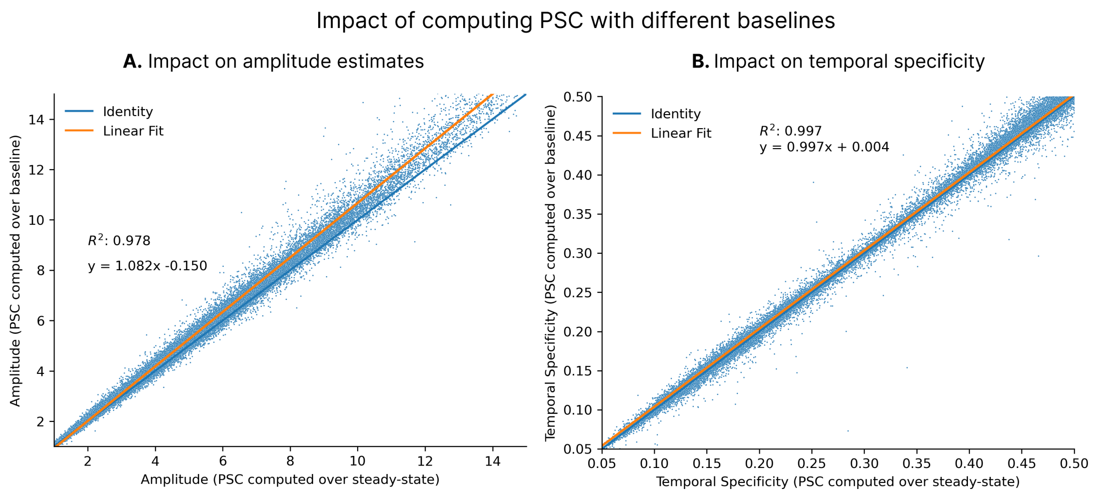


***Supplementary Figure 4. The impact of baseline choice on PSC estimates and on temporal specificity.*** (A) The response amplitude (as PSC) estimated using as baseline the mean of the steady-state response (horizontal axis) plotted against the response amplitude (as PSC) estimated using as baseline the mean of the 10-s period of the acquisition before onset of stimulation (vertical axis). (B) Analogous to panel A, except temporal specificity is plotted instead of amplitudes. Each dot represents a single voxel. A linear fit indicates that amplitudes are approximately 8% larger using the initial period as a baseline for PSC computation ($R^2$=0.978, panel A). Almost no difference is seen for the temporal specificity estimates, with a comparison between values obtained with both approaches showing strong agreement ($R^2$=0.997, panel B). Voxelwise variance is likely influenced by the noise in the estimates of the mean in the baseline period since the mean is computed over a short period with few datapoints. The choice of PSC baseline therefore does not affect results on temporal specificity.

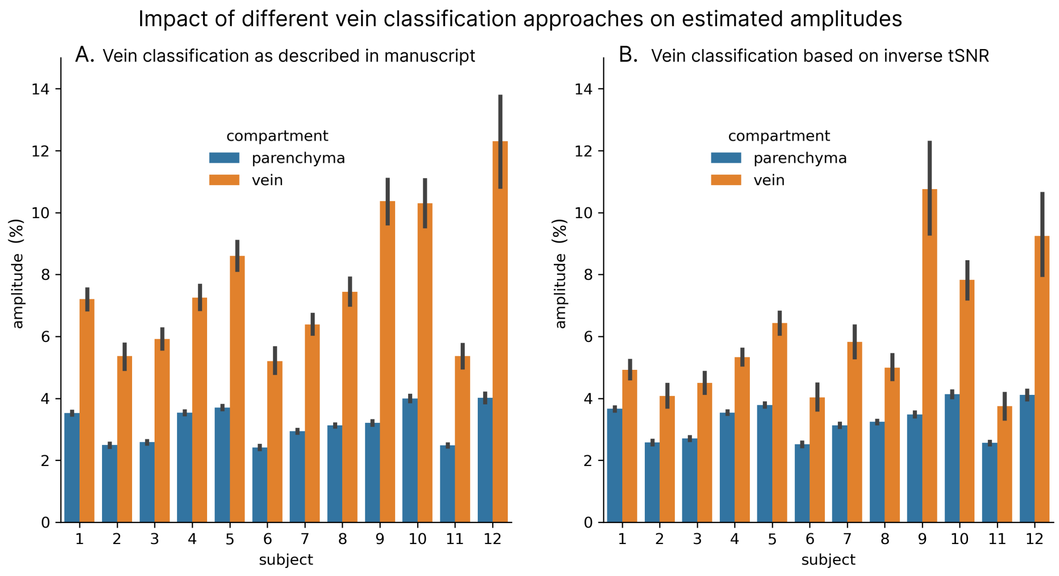


***Supplementary Figure 5. Impact of venous classification approach on estimated amplitudes at 0.10 Hz.*** (A) Bar plot showing mean amplitude for each subject across compartments as computed in the current study. (B) Analogous to the bar plot in panel A, but with the venous compartment computed using inverse tSNR as a metric. Inverse tSNR may be misclassifying low-amplitude parenchymal voxels as veins, thus reducing the mean amplitude of the "vein" class, even if inverse tSNR correctly identifies ascending intracortical venules.

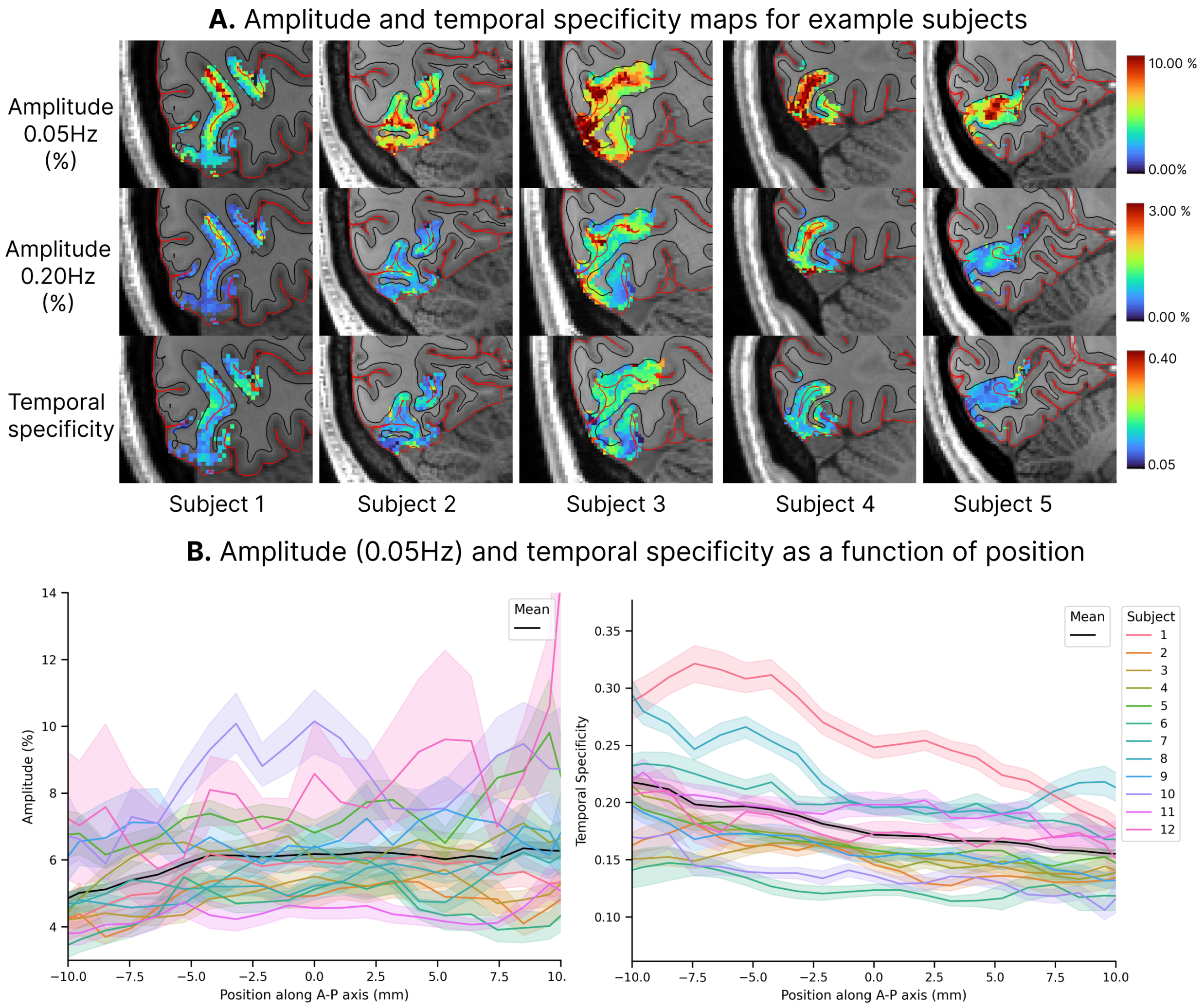


***Supplementary Figure 6. Demonstration of variability of amplitude and temporal specificity across the A–P axis across subjects.*** (A) Amplitude response and temporal specificity maps at the single-subject level shown for 5 out of 12 subjects. Maps are shown overlaid on anatomical space, with the FOV placed over V1. Cortical surfaces are shown in black (white matter surface) and red (pial surface). Top row: Estimated amplitude responses to the 0.05-Hz task. Middle row: Estimated amplitude responses for the 0.20-Hz stimulus. Amplitudes are clearly dominated by the cortical depth effect, and show substantial spatial heterogeneity Bottom row: Temporal specificity, computed as the ratio of the 0.20-Hz by the 0.05-Hz amplitudes. For temporal specificity maps, however, no cortical depth dependency is apparent, and maps are much more sensitive to position across V1. (B) Amplitude and temporal specificity as a function of position across all 12 subjects. Shaded areas represent 95% confidence interval of the mean, computed over voxels. Substantial variation can be seen across subjects, nonetheless, for all subjects a trend of temporal specificity increasing towards anterior V1 is apparent.

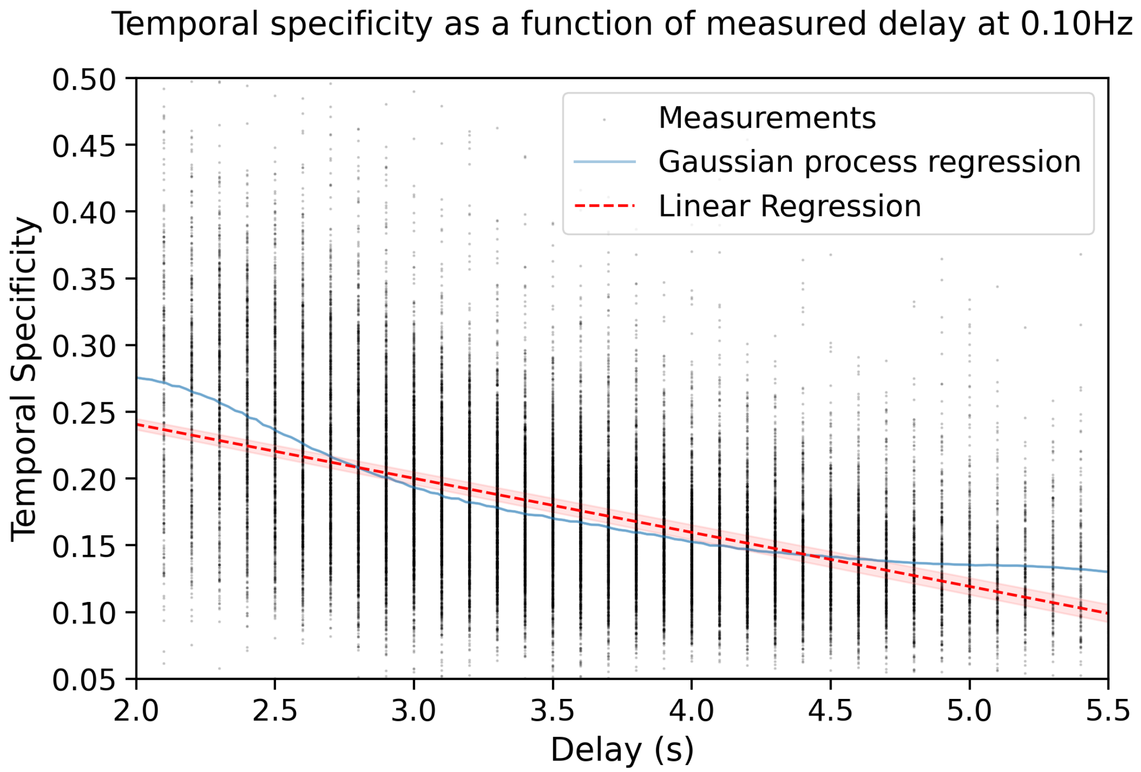


***Supplementary Figure 7. A direct relationship between estimated delays and the temporal specificity.*** Plot of temporal specificity as a function of delay. Each dot represents a voxel and data is shown for all subjects. Red line represents a linear regression (shaded bars representing 95% confidence interval), blue line a gaussian process regression. We use the GP regression only to illustrate that the relationship between delays and temporal specificity is not strictly linear. At delays larger than about 3.5 s, there is a smaller impact to the temporal specificity than at earlier delays, but the relationship exhibits a strict monotonic decrease with increasing delays, and delays are linked with temporal specificity.

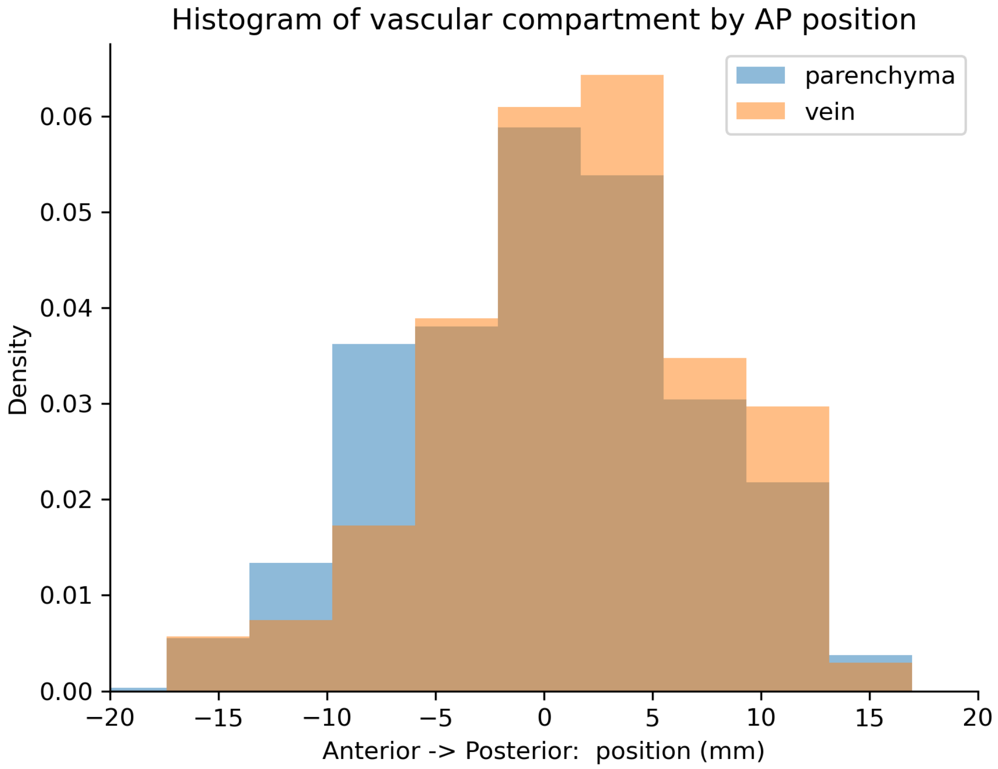


***Supplementary Figure 8. Histogram of the normalized density of venous and parenchymal voxels across the anterior-posterior axis.*** There is a slight sampling bias in that a higher density of voxels detected as veins is seen in posterior V1. Our data, however, cannot unambiguously tell us whether indeed more veins are present in posterior V1. Histograms were generated from data pooled from all subjects. Density is normalized such that integral is equal to 1.

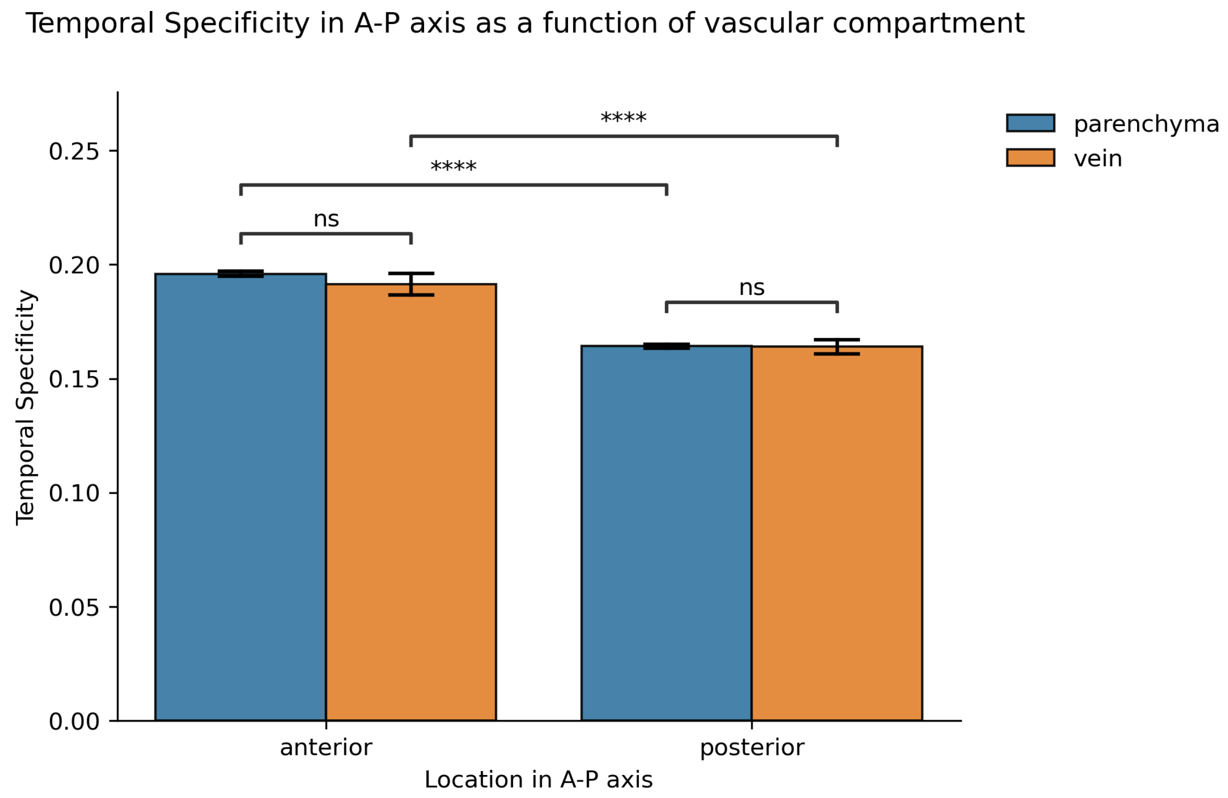


***Supplementary Figure 9. Temporal specificity stratified by compartment and position along the A–P axis of V1.*** Temporal specificity is significantly higher in anterior V1 in both venous and parenchymal compartments. Notably, veins are also affected by position, such that veins in the posterior V1 also have lower temporal specificity than veins in anterior V1. No significant differences are found in temporal specificity in either anterior or posterior V1. False negatives or positives in vein classification are unlikely to influence our positional findings in temporal specificity—undetected veins in anterior V1 would likely reduce temporal specificity, thus if they were properly detected and voxels properly classified we would expect an even higher temporal specificity in parenchymal voxels within anterior V1. Stars indicate high statistical significance, $p < 10^{-5}$ (Mann-Whitney-Wilcoxon test, two-sided with Bonferroni correction).
